# In vivo metabolite tracing of T cells

**DOI:** 10.1101/2020.08.21.261503

**Authors:** Ryan D. Sheldon, Eric H. Ma, Lisa M. DeCamp, Kelsey S. Williams, Russell G. Jones

**Affiliations:** Metabolic and Nutritional Programming, Center for Cancer and Cell Biology, Van Andel Institute, Grand Rapids, MI 49503, USA

**Keywords:** stable isotope tracing, immunity, metabolism, immunometabolism, T cells, cell sorting

## Abstract

T cells are integral players in the adaptive immune system that readily adapt their metabolism to meet their energetic and biosynthetic needs. A major hurdle to understand physiologic T cell metabolism has been the differences between in vitro cell culture conditions and the complex in vivo milieu. To address this, we have developed a protocol that merges traditional immunology infection models with whole-body metabolite infusion and mass spectrometry (MS)-based metabolomic profiling to assess T cell metabolism *in vivo*. In this protocol, pathogen-infected mice are infused via the tail vein with an isotopically labeled metabolite, followed by rapid magnetic bead isolation to purify T cell populations and then stable isotope labeling (SIL) analysis conducted by MS. This procedure enables researchers to evaluate metabolic substrate utilization by specific T cell subpopulations in the context of physiological immune responses in vivo.

## Introduction

Cellular metabolism underlies the essential functions of all cell types, providing building blocks for growth, redox balance, and energy. It is widely appreciated that metabolic dysregulation drives or contributes to some illnesses, such as diabetes and cancer; thus, a deeper understanding of metabolism can provide insight into the biological processes that underlie both normal physiological responses and pathophysiology, potentially uncovering new therapeutic options. A key technique used in the study of metabolism is stable isotope tracing, a process which involves tracking “heavy labeled” elements such as carbon (C) or nitrogen (N) through biochemical pathways in living cells or tissues. This technique has elucidated key metabolic pathways engaged by cells and provided insight into the usage of metabolites—like glucose^1,2^, glutamine^3^, acetate^4^, and serine^5^—to fuel specific cellular processes.

It is becoming clear that the metabolic profiles of cells are tissue- and context-dependent^6,7^, and cellular surroundings play an important role in shaping a cell’s dynamic metabolic network. While stable isotope tracing has highlighted diverse metabolic pathways that can support cell function, these studies are often done *in vitro* using culture media designed to maximize in vitro proliferation. As a consequence, in vitro culture conditions often do not accurately recapitulate in vivo environmental conditions or metabolic processes^8^. For example, glutamine is a major anaplerotic carbon source for lung tumor cells in vitro but not in vivo^6^. To address this gap, recent efforts have demonstrated the utility of media designed to mimic small-molecule concentrations in human plasma^9,10^. However, other factors such as fluid dynamics, oxygen tension, cytokines, and complex intracellular interactions make it difficult to replicate the in vivo milieu—and therefore in vivo metabolism—in cell culture. As an alternative strategy, many labs have turned to using in vivo tracing approaches to understand metabolic dynamics in vivo.

Recent advances in implementing stable isotope tracers in vivo have led to a more accurate understanding of cellular metabolism in a physiological context. This has been particularly true in the field of cancer metabolism, where stable isotope tracers have been successfully used to assess cancer metabolism in vivo through tumor xenografts or solid tumors ^6,11–14^. Part of this success is due to the relative ease of dissecting tumors and quickly quenching metabolism by flash freezing in liquid nitrogen after in vivo tracer administration. A guiding principle for metabolomics studies is to quench metabolism as rapidly as possible at the end of the experimental period. Thus, an accurate snapshot of in vivo tumor metabolism can be captured. This approach is limited, however, in its ability to discern between cell subtypes, such as is needed in immunological studies. For example, in order to characterize the in vivo metabolism of specific immune cell subtypes such as T cells, it is necessary to rapidly isolate the desired cell population to minimize changes in metabolism during processing, a goal at odds with time-intensive traditional cell sorting procedures.

T cells are integral players in the adaptive immune system, tuning antigen-specific immunity and developing immunological memory. These functions are accomplished by specialized cellular sub-populations with unique functions and corresponding metabolic demands. Upon immunological challenge, quiescent naïve T cells (Tn) rapidly differentiate into highly proliferative effector T cells (Teff). This transformation requires a similarly rapid metabolic adaptation to provide the energy and small-molecule building blocks (e.g., nucleotides) needed to support growth. Following pathogen clearance, Teff cells employ a different set of metabolic pathways in order to transition to memory T cells (Tmem), quiescent T cells with rapid recall capacity. These various metabolic pathways may be therapeutically targeted to stimulate or suppress T cell function, promoting protective immunity or diminishing autoimmunity, respectively. However, a full understanding of these targetable metabolic pathways has remained elusive due to a lack of methods that characterize metabolism in a physiologic context.

T cell subpopulations can be purified using fluorescence-activated cell sorting (FACS), but the process is lengthy; fluorochrome staining followed by flow cytometry-based sorting can take hours to accumulate sufficient cells for detection via mass spectrometry. During this process, cells are subjected to sorting conditions, which are much different than the in vivo milieu, allowing time for metabolite exchange with the sorting buffers. As such, the post-sort metabolome is much different than the state in which it was collected ^15,16^. Here, we have developed a workflow to interrogate the in vivo metabolism of T cell subpopulations by coupling in vivo stable isotope tracer infusion to rapid bead-based cell isolation.

### Development of the protocol

A major challenge to tracing T cell metabolism in vivo is having to quickly isolate the T cell population and quench metabolism following an in vivo infusion. Where traditional protocols take 1–1.5 hours to isolate cell types, we wanted to develop a method that is faster and less stressful to the T cells in order to preserve in vivo labeling patterns. Such a strategy has been successfully applied to metabolomics of mitochondria, where genetically tagged mitochondria can be isolated in ~15 minutes ^17^. Along these lines, we developed a magnetic bead-based Thy1.1^+^ cell isolation method, which is able to isolate Thy1.1^+^ T cells in 35 minutes without affecting the label incorporation of most metabolites^18^.

Our bead-based isolation method dramatically decreases processing time and cell stress when compared to traditional FACS approaches. Like FACS however we also observe increased variability and general decreases in the abundance of many metabolites from our bead sorting procedure. This is likely due to metabolite exchange with the bead sorting buffer, as metabolomic analysis of the sorting buffer displayed significant labeled metabolites present at the end of the sorting protocol. Therefore, this method may be quantitatively unreliable for reflecting in vivo metabolite abundances. However, the mass isotopologue distribution (MID; see Box 1), which represents the fraction of a given metabolite that contains a certain labeling pattern, is unaffected by processing variability for most metabolites. Therefore, the current method analyzed MID patterns to qualitatively understand metabolic pathways in vivo.

In order to study the metabolism of CD8^+^ T cells over the course of activation/infection, we utilized OT-I transgenic T cells. OT-I T cells express an H-2D^b^-restricted T cell receptor (TCR) specific for the ovalbumin (OVA) epitope (residues 257–264), allowing for specific T cell activation via OVA peptides. In vivo, OT-I T cell activation is achieved by adoptively transferring OT-I T cells into host animals (C57BL/6), followed by infection with *Listeria monocytogenes* modified to express ovalbumin antigen (*LmOVA*). This allows for the specific activation of a large amount of OT-I T cells in vivo to generate sufficient cell numbers for metabolomic studies on the response of CD8^+^ T cells to antigen.

The use of OT-I T cells that genetically express the CD90.1 (Thy1.1) cell surface protein allows us to differentiate the adoptively transferred OT-I T cells from endogenous T cells from the host C57BL/6 mice, which express the CD90.2 (Thy1.2) cell surface protein. This differentiation is crucial, as we utilize a positive-selection bead isolation kit that will purify Thy1.1 cells from the mice, allowing us to rapidly isolate the antigen specific OT-I T cells at desired time points during infection to probe the metabolism of CD8^+^ T cells in response to antigen. This method is amenable for sorting T cells or other immune cells using antibodies that target abundant endogenous (or transgenically-expressed) cell surface proteins.

##### Box 1 on nomenclature

In stable isotope labeling (SIL) metabolomics, the terms “isotopologue” and “isotopomer” are often used interchangeably to describe a given labeled species of metabolite with “n” number of labeled carbons. In fact, “isotopologue” refers to a metabolite with a specific number of incorporated labeled carbons, irrespective of the position of the carbons, whereas “isotopomer” refers to a labeled metabolite with a specific positional arrangement of the labeled carbons. For example, in a 3-carbon molecule, there are four possible ^13^C isotopologues (M+0, M+1, M+2, and M+3) but eight possible isotopomers (See **Figure B1** below). In most mass spectrometry (MS)-based SIL studies, positional labeling is not determined and *isotopologues* are the measured outcome. Isotopomers are determined when positional labeling is elucidated through compound fragmentation patterns in MS experiments or by nuclear magnetic resonance (NMR).

**Figure B1.**
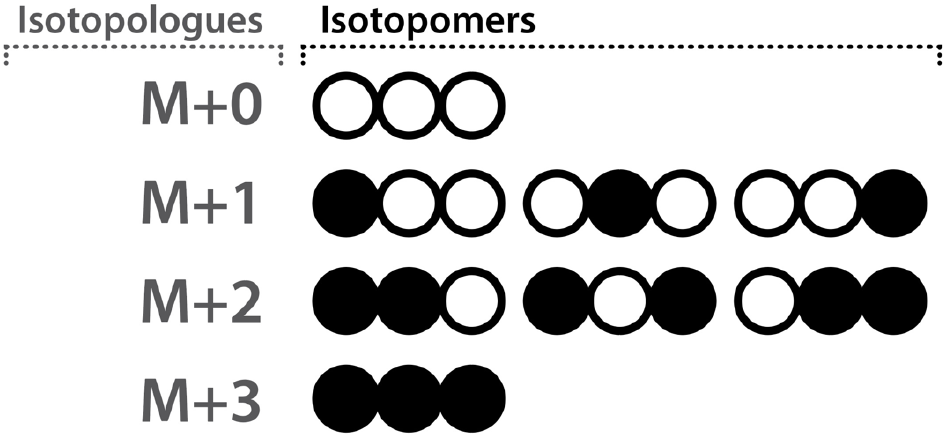
The isotopomers and isotopologues possible for a 3-carbon molecule. (to be included inside the box)

### Applications of the method

The in vivo cellular phenotype is determined by complex environmental interactions that are severely perturbed or misrepresented during standard culture. Here we describe a workflow that couples stable isotope infusion with rapid cell isolation in order to systematically interrogate the in vivo metabolic pathways of T cells, specifically over the course of infection. Our recent work used this method to identify key differences in T cell metabolism in culture versus in vivo^18^. Although in traditional culture conditions, Teff cells exhibit classical Warburg metabolism characterized by high aerobic glycolysis and lactate production, we demonstrated that Teff cells activated in vivo and isolated with the workflow described herein display markedly lower glucose flux to lactate. Instead, these cells rely more heavily on oxidative phosphorylation for ATP production and utilize glucose for nucleotide biosynthesis. These data highlight the importance of evaluating metabolism in vivo, and this protocol can be easily adapted to the study of other tissues and/or cell types across a range of experimental conditions.

### Comparison with other approaches

There are multiple approaches to *in vivo* stable isotope tracing that are compatible with downstream T cell isolation. Here we discuss the advantages and disadvantages of each, and a summary is presented in **Table 1**.

**Table 1.** Advantages and disadvantages of common in vivo labeling approaches.

| Method of isotope administration | Advantages | Disadvantages |
| --- | --- | --- |
| Single/multiple bolus (IP/IV) | - Simple<br>- Conscious animal | - Not steady state<br>- Nutrient spikes affect physiology |
| Bolus followed by continuous tail vein IV infusion, sedated | - Steady state labeling<br>- Less complicated than jugular catheterization | - Anesthetic-specific effects on metabolome<br>- May slightly elevate circulating levels of infused compound |
| Bolus followed by continuous infusion via jugular vein catheter | - Conscious animal<br>- Steady state labeling | - Requires surgery, specialized expertise and equipment |
| Bolus (gavage) | - Simple<br>- Conscious animal<br>- Accounts for digestion/absorption | - Not steady state<br>- Influenced by digestion/absorption |
| Diet or drinking water | - Simple<br>- Conscious animal<br>- Potentially steady state<br>- Accounts for digestion/absorption | - Variability due to intermittent hydration and feeding behavior<br>- Taste effects (e.g., glutamine is bitter, glucose is sweet)<br>- Low dosage control<br>- Influenced by digestion/absorption |

#### 1. Single or multiple boluses via tail vein intravenous (IV) injection, intraperitoneal (IP) injection, or gavage

One of the simplest approaches to in vivo isotope tracing is the discrete bolus injection of tracers. This has the advantage of avoiding the use of anesthesia and relatively simple to perform. However, instead of achieving constant tracer enrichment in the bloodstream, multiple discrete injections causes repeated peaks in tracer nutrient levels in the blood stream. This can have unnatural systemic metabolic effects, such as glucose spikes causing increases in insulin secretion with each injection. Downstream data analysis also becomes more complex. At any given time, the cells are exposed to a varying level of tracer, and data analysis must account for the tracer turnover rate.

#### 2. Bolus followed by continuous infusion via tail vein

The first administration of a bolus allows for the initial flooding of the system with heavy-labeled metabolite, improving the probability of cellular uptake of heavy-labeled metabolite over the naturally occurring metabolite in circulation. Following the bolus, the tracer is continuously administered at a set rate into circulation via the tail vein until the experimental end point. This approach allows the system to reach a metabolic steady state (i.e., the rate of metabolite production and clearance are equal, so there are no net changes in concentration). This is critical because the labelling of endogenous metabolites is directly proportional to the level of labelling achieved in the infused metabolite. Since infused tracer levels in circulation are at steady state, the tracer will permeate into tissue metabolic pathways at a proportional rate. Thus, steady-state labeling of entire pathways can be evaluated.

However, like with most techniques, there are disadvantages that should be considered and noted. First, the administration of the tracer in a bolus/infusion increases circulating levels of the metabolite, artificially presenting the cells with a transiently higher concentration than typically observed in vivo. Second, for continuous tail vein infusion (2–6 hours), mice must be sedated, typically with ketamine or isoflurane, for the duration of the infusion. Anesthetics can affect metabolism and should be selected based on compatibility with desired metabolic outcomes. This is discussed further in “Limitations”.

#### 3. Bolus followed by infusion via jugular vein catheter

Infusion via surgical catheterization of the jugular vein shares multiple advantages and disadvantages with infusion by the tail vein. Infusion through the jugular vein has the advantage of not requiring the mice to be sedated during the length of the infusion, which removes the potential concerns of sedative influencing cell metabolism, negative drug interactions, and mice being immobile during the length of the infusion.

Some disadvantages associated with jugular vein infusions are the specialized expertise needed to perform surgery to insert the catheter into the jugular vein, and the recovery time required prior to experimental use of the catheter. There is also the added cost associated with having to house mice individually in separate cages.

#### 4. Labeled nutrient in diet or drinking water

A more passive approach to *in vivo* tracing is through oral administration of the tracer by incorporating the tracer into the drinking water or diet. Introducing the tracer through the animal diet has the advantage of modeling nutrition-based metabolite incorporation into DNA, proteins, and lipids, as the tracer is digested, absorbed, and introduced into circulation at normal levels over time, keeping the mice physiology normal. This approach has a similar disadvantage as multiple injections of a bolus of tracer, in that circulating tracer nutrient levels ebb and flow, albeit less dramatically.

#### Additional considerations for in vivo tracing, irrespective of method

There are additional considerations that need be taken into account, irrespective of the chosen method for stable isotope tracer administration. First, the length of fasting prior to tracer administration is critical. Prior to in vivo tracing, mice are typically fasted to achieve a consistent metabolic baseline and decrease the influence of dietary nutrients. Detailed information should be reported describing the duration of fasting (6 hours–overnight) as well as the time of day the fast is done. This improves reproducibility, as multiple studies have shown the close relationship between metabolism and the circadian clock^19–21^. Secondly, the metabolome is easily perturbed by stress. Care should be taken to handle the mice as minimally and consistently as possible during all phases of the experiment.

### Expertise needed

Listeria monocytogenes is a Biosafety Level 2 (BSL-2) pathogen. Adoptive transfer and *LmOVA* infection techniques can be conducted by laboratory staff with basic animal husbandry expertise, tissue culture training, and BSL-2 containment protocols. Tail vein infusions can become routine but require considerable practice. In our lab, practicing on 5–10 mice per day for three weeks is usually sufficient to fully train new personnel in the technique. Personnel are considered proficient when they can successfully infuse 10 mice in a row. Trained personnel have an 85%–95% success rate with this technique.

SIL analysis requires expensive instruments and specialized training. Most often, this aspect of the workflow can be completed in a metabolomics core facility or in collaboration with a lab that has mass spectrometry equipment and expertise.

### Limitations

Investigating the in vivo metabolic state of specific cell types is difficult because the conditions and length of processing can dramatically affect the metabolome. The protocol we detail here can largely alleviate this difficulty through rapid magnetic bead sorting. Though this method is much shorter than the flow-cytometry alternative, there is potential for metabolism to continue during this time. Accordingly, the following limitations should be considered:

1. Consistency in timing between replicates is critical to minimize inter-sample variability. This contrasts with the “moving as fast as possible” approach for each replicate. Each step should be practiced, timed, and a schedule established and followed.
2. Using pre-chilled buffers and keeping samples on ice or in refrigerated centrifuges will slow metabolism and limit processing artifacts.
3. Total pool sizes of metabolites in sorted T cells may be unreliable. During processing there will be loss of intracellular metabolites due to exchange with sorting medium. The labeled fraction of most metabolites, however, is unaffected by this exchange. Therefore, this method allows interrogation of metabolic pathways active in different in vivo conditions by examining *labeling patterns*.
4. Investigators should be mindful of effects of the sorting process on cell physiology that may alter metabolic pathways. While we have not observed this directly, it must be considered when interpreting results.
5. Additional control and validation experiments should follow important findings. Purity and activation phenotype of isolated cells should be validated using established methods, such as flow cytometry. Metabolite labeling patterns in vivo should be validated with secondary methods such as ex vivo SIL or Seahorse analysis.

Another potential limitation is that this protocol uses long-term (2.5–6 hours) isoflurane anesthesia during the sorting process. This obviates the need for complicated surgical approaches otherwise necessary to reach metabolic steady state. While isoflurane has been shown to be minimally disruptive to the metabolome^22^, it can alter glucose metabolism through inhibition of mitochondrial oxidative phosphorylation^23^. Further, isoflurane and other common anesthetics such as ketamine can activate xenobiotic metabolic pathways in the liver^24,25^, which could be particularly relevant to pharmacological investigations. Considering this, the metabolic effects of isoflurane or any anesthetic approach should be considered in the context of the specific goals of the investigation.

### Experimental Design

#### Overview

Here we provide a step-by-step procedure to assess metabolic stable isotope labeling patterns in T cells in vivo in the context of Listeria infection (**Fig. 1**).

**Figure 1.**
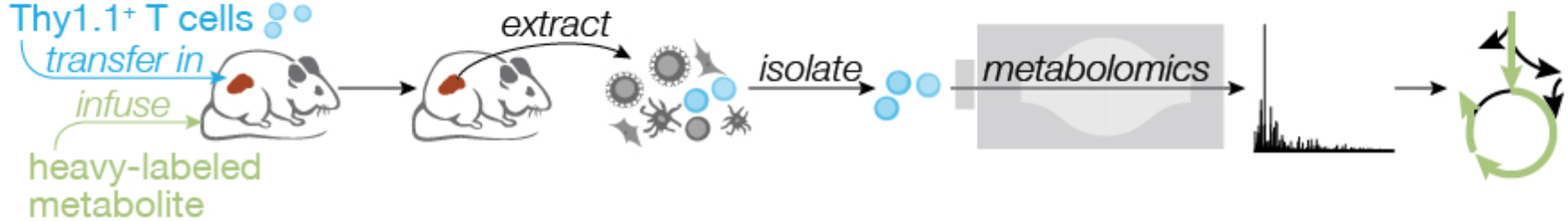
Overview of the procedure. Schematic of workflow, with graphs of data at critical steps.

#### Adoptive transfer and Listeria infection

CD8^+^ T cells are isolated from OT-I Thy1.1^+^ mice and are transferred into C57BL/6 mice. The C57BL/6 mice are then infected with *L. monocytogenes* expressing the OVA epitope (LmOVA).

#### In vivo perfusions

Two to six days post-infection, anesthetized mice (animal feed removed 6 hours prior to tail vein infusion) are perfused via tail vein with a labeled metabolic substrate of interest for 2.5–6 hours.

#### T cell isolation

Following labeled metabolite perfusion, T cells are isolated from spleen homogenates. Magnetic bead sorting allows for rapid separation of Teff cells (or Tn cells if desired) to capture in vivo metabolite labeling in each subpopulation.

#### SIL Metabolomics

Metabolites are extracted from frozen T cells and analyzed by liquid chromatography-mass spectrometry (LC/MS) or gas chromatography-mass spectrometry (GC/MS) for SIL.

## Materials

### Reagents

- C57BL/6 mice
- Thy1.1^+^OT-I^+^ mice (C57NL/6-Tg(TcraTcrb)1100Mjb/Crl crossed to B6.PL-*Thy1^a^*/CyJ)
- Isoflurane
- Compressed oxygen
- HBSS (Cat# 311-510-CL, WisentBioProducts)
- Easy-Sep media (PBS with 2%FBS and 1mM EDTA)
- EasySep™ Mouse CD8^+^ T cell isolation kit (Cat# 19853A, STEMCELL Technologies)
- EasySep™ Mouse CD90.1 Positive Selection kit (Cat# 18958, STEMCELL Technologies)
- EasySep™ Mouse Naïve CD8+ T cell Isolation Kit (Cat# 19858, STEMCELL Technologies)
- FACS antibodies (See Table 3)
- LC/MS grade methanol (Cat# A456, Fisher Scientific)
- LC/MS grade acetonitrile (Cat# A955, Fisher Scientific)
- LC/MS grade water
- LC/MS grade ammonium hydroxide (Cat# A470, Fisher Scientific)
- LC/MS grade ammonium acetate (Cat# A11450, Fisher Scientific)
- Medronic acid (Cat# 5191-3940, Agilent Technologies)
- D27 myristic acid (Cat# 366889, Sigma)
- MTBSFA+1%TMCS (Cat# 00942, Sigma)
- Pyridine (Cat# PX2012, Sigma)
- Methoxyamine HCl (Cat# 89803, Sigma)
- Stable isotope tracers (see **Table 2**)

**Table 2.** Concentrations, infusion rates, and expected enrichment of common tracers for a 20–25 g mouse

| Tracer | Concentration (mg/mL) | Bolus dose (μL) | Infusion rate (μL/min) | Expected blood enrichment (%) | Catalog # |
| --- | --- | --- | --- | --- | --- |
| U-[<sup>13</sup>C]-Glucose | 100 | 120 | 2.5 | 60–80% | CLM-1396 |
| U-[<sup>13</sup>C]-Glutamine | 40 | 150 | 3.0 | 50–60% | CLM-1822 |
| U-[<sup>13</sup>C]-Acetate | 40 | 150 | 3.0 | Unknown. Blood citrate ~20% | CLM-440 |
| U-[<sup>13</sup>C]-Serine | 15 | 150 | 3.0 | 50–60% | CLM-1075 |

**Table Footnote.** Table is based on work by several laboratories<sup>6,27–30</sup>. Provided catalogue numbers are from Cambridge Isotope Laboratories, Tewksbury, MA , USA.

**Table 3.** Antibodies for Flow Cytometry.

| Fluorochrome | Surface Protein | Dilution | Catalog # |
| --- | --- | --- | --- |
| FITC | KLRG1 | 1/300 | (ThermoFisher) 11-5893-82 |
| PE | CD8 | 1/300 | (ThermoFisher) 12-0081-85 |
| PECy7 | CD44 | 1/300 | (ThermoFisher) 25-0441-82 |
| APC | Thy1.1 | 1/300 | (ThermoFisher) 17-0900-82 |
| APCCy7 | Viability | 1/1000 | (ThermoFisher) 65-0865-14 |
| BV421 | CD127 | 1/100 | (Biolegend) 135024 |
| BV605 | CD4 | 1/300 | (Biolegend) 100548 |

### Equipment

- Heating mat
- Laboratory tape
- Paper towels
- 27G scalp vein assembly (Cat# 26709, EXELint)
- Plastic tubing (Cat# MRE-033, Braintree Scientific)
- 25G needle (Cat# 305127, BD Biosciences)
- 1 mL syringes
- 50 mL Falcon tubes
- 29G insulin syringes (Cat# 329420, BD Biosciences,)
- EasyEights™ EastSep™ Magnet (Cat# 18103, STEMCELL Technologies)
- EasySep™ Magnet (Cat# 18000, STEMCELL Technologies)
- Autosampler vials with fused insert (Cat# 29391-U, Sigma)
- 70 μm cell strainer (Cat# 352350, Corning)
- Falcon round-bottom polystyrene tubes 5 mL (Cat# 14-959-5, Fisher Scientific)
- HPLC Column (SeQuant ZIC-pHILIC 5 μm polymer 150 x 2.1 mm, Cat#150460, Millipore Sigma)
- GC column (J&W DB-5ms GC Column with 10m DuraGuard, Cat# 122-5532G, Agilent Technologies)
- Syringe pump
- Isoflurane vaporizer
- Vortex
- Benchtop centrifuge/microcentrifuge
- Flow cytometer
- Heat block
- Speed vac
- Thermo Vanquish Horizon HPLC
- Thermo Orbitrap ID-X mass spectrometer with electrospray ionization (ESI)
- Agilent 7890 gas chromatograph system
- Agilent 5977b mass selective detector (MSD) with electron impact (EI) ionization

## Procedure

### Thy1.1^+^OT-I^+^ T cell adoptive transfer and listeria infection. Timing: 2 days

#### Day 0

1. Isolate CD8^+^ T cells from the spleen of Thy1.1^+^OT-I^+^ mice via negative selection using EasySep™ Mouse CD8^+^ T cell isolation kit following manufacturer’s protocol.
2. Re-suspend isolated Thy1.1^+^OT-I^+^ CD8^+^ T cells in HBSS to desired concentration for injection into C57/BL6 mice via tail vein in 200 μL

a. For OT-I Thy1.1^+^CD8^+^ at 2.5 days post infection (dpi) inject 2.5 × 10^6^ cells/mouse.
b. For OT-I Thy1.1^+^CD8^+^ at 6 dpi, inject 5 × 10^4^ cells/mouse. *Note:* Injection of too few cells can impair sufficient recovery at early time points, and too many cells can alter the infection trajectory (Ref: PMID 1755991). The number of cells provided here is based on previous work^26^, serves as a guide, and should be determined empirically in each lab.

#### Day 1

3. Grow attenuated listeria monocytogenes expressing the OVA epitope for infection of C57BL6 adoptively transferred with Thy1.1^+^OT-I^+^ CD8^+^ T cells
4. Infect at 2 × 10^6^ CFU/mouse by IV injection (200 μL/mouse of 1 × 10^7^ CFU/mL).

### Preparing infusion lines. Timing: 30 minutes

1. From the 27G scalp vein assembly (EXELint, 26709), cut and strip the attached line assembly from the base of the butterfly wings, exposing the metal back end of the needle. Cut one of the wings off to reduce the weight (**Figure S1A**).
2. Cut a length of tubing (Braintree Scientific, MRE-033) about the width of the infusion heating pad (18–20”).
3. Attach a 25G (blue) needle (BD #305127) to a 1 mL syringe. Poke the needle into a Styrofoam box, and cut the needle so that the metal end is about half an inch long (**Figure S1B**). Trim it down about a quarter inch over a disposal container. Inspect the end of the now-blunted needle to ensure that it is not collapsed. A small rounded hole should be visible. If not, cut of a small amount and check again.
4. Glide the tubing over both the blunted end of the 25G needle (attached to the syringe) and the 27.5G scalp vein needle (**Figure S1C**).

### Preparing tracers. Timing: 30 minutes

1. Weigh out the tracer into a tared 50 mL tube.
2. Dissolve tracer in the appropriate volume of sterile saline (9 g/L NaCl) to achieve desired concentration. Recommended concentrations are based on previously published reports^6,27–30^. *Important*: do not use PBS or other buffered solutions as these can cause ion suppression.
3. *Note*: Concentrations and infusion rates are for a 20–25 g mouse. If mice are of different weights, especially if weight differences are expected between groups, then concentration and/or infusion rate can be modified to make tracer load/g of mouse consistent. If differences in body composition are expected, it is suggested to base dosing on lean mass (measured by NMR/echo-MRI).

### Preparing syringes. Timing: 30 minutes

*Important:* these steps are critical to remove all air from the infusate. An air embolism can increase incidence of mortality.

1. Using a 1 mL syringe without a needle, draw up roughly 400 μL of tracer. Then invert the needle and pull back a few hundred μL of air. This is to prevent volume loss when flicking the syringe. Flick the syringe until no bubbles are observed and all air is at the top.
2. Invert the needle and expel the air back into the falcon tube.
3. Now that the syringe contains solution and no air, put the end of the syringe back into the solution, and retract to about the 1 mL mark. Inspect to make sure that there are no bubbles in the syringe.
4. Attach the infusion line.
5. Put the end of the infusion line into the tube, and flush the air from the line, bringing the plunger on the syringe to the 700 μL position.
6. Check the line and syringe again to make sure there are no bubbles. If you see a bubble, flush everything back into the falcon tube, and start again.

### Sedating and preparing the mouse. Timing: 15 minutes

*Overview*: Mice need to remain sedated throughout the duration of tracer infusion (2–6 hours). We prefer isoflurane delivered with oxygen due to ease of delivery and precise dosage control. An alternative approach using a sedation cocktail is also presented below.

*Note*: Anesthesia, regardless of type, can influence metabolomic readouts. Therefore, it is important to understand the minimal dose required to maintain a constant anesthetic plane and ensure that this dosage is consistent across replicates in an experiment.

*Preferred Method*: Using an isoflurane sedation method.

1. Place the mouse in a drop box perfused with isoflurane and oxygen using an isoflurane vaporizer.
2. Transfer the mouse to the heating pad once it is unable to walk (~2 min). Place the nose of the mouse in a nosecone to continue O2/isoflurane delivery. *Note*: It is important to consistently use the minimal effective dose of isoflurane. As a general guideline for initial anesthesia, the mouse should be immobile after ~2 minutes in the drop box. When on the nosecone, the proper dose can be estimated by removing the nosecone from the mouse. If the proper dosage is being used, then the mouse should begin to stir within a minute. *Important*: The mouse should be monitored continuously while sedated. If the mouse begins to move, make small incremental increases in the isoflurane dose. Increases in respiration rate or change in skin color can indicate overdose of isoflurane.
3. Once fully sedated, use a razor blade to shave the insides of the legs so that the femoral vein is visible. This helps for doing blood draws during the experiment.
4. Tape the mouse supine to the heating pad by its paws and tail (after infusion needle is correctly inserted). *Important:* Taping the tail is necessary to prevent the mouse from disturbing the tail-vein needle if it begins to wake.

*Alternative Method:* using a sedation cocktail of ketamine, xylazine, and acepromazine

*Overview:* Sedation is accomplished by giving an initial bolus of sedation cocktail to achieve a steady anesthetic plane for 1–1.5 hours, followed by smaller maintenance doses to achieve sedation for the entire experiment (4 hrs).

*Important:* Overdosing or being generous with the amount injected will kill the animal. Especially when using a new preparation, start on the low end of the dosing schedule. The doses recommended here are on the low-end and should not risk death in an otherwise healthy animal.

*Important*: The cocktail will degrade on light exposure.

*Important:* The mouse should remain on the heating pad throughout sedation to prevent hypothermia and cause dilation of the tail vein.

1. Prepare sedation cocktail of Ketamine (50 mg/mL), Xylazine (50 mg/mL) and acepromazine (2 mg/mL).
2. Load 29G insulin syringes (BD #329420) with the exact amount needed ahead of time (and store in a drawer or other light-protected space).
3. Dosage:

a. For mice that weigh 15–30 g

i. Initial dose: 50 μL
ii. Maintenance dose: 10 μL
b. For mice that weigh > 30 g

i. Initial dose: 60 μL
ii. Maintenance dose: 15 μL
4. Set a 5-minute countdown timer.
5. Give the mouse the initial intraperitoneal (IP) injection of sedation cocktail.
6. Start the timer.
7. If the dosage is correct, the mouse will begin to wobble after the first minute, it will be unable to walk by the second minute, and most of the muscular tremors should stop by the fourth or fifth minute.
8. Transfer the mouse to the heating pad once it is unable to walk (~2 min).
9. Once fully sedated, use a razor blade to shave the insides of the legs so that the femoral vein is visible. This helps for doing blood draws during the experiment.
10. Tape the mouse supine to the heating pad by its paws and tail (after infusion needle correctly inserted). *Important:* Taping the tail is necessary to prevent the mouse from disturbing the tail-vein needle if it begins to wake.
11. Continuously monitor the mouse for signs of waking throughout the procedure. In the minutes before a mouse begins to wake, they tend to urinate, although this is not always the case. The first clear sign that a mouse is waking is that its nostrils will repeatedly flare as the mouse begins sniffing. If the mouse has long whiskers, this is very obvious since they will start to fan.
12. When the mouse begins to wake, deliver a maintenance dose of sedation cocktail immediately. The maintenance dose (described in Step 3 of this section) should result in an additional 45–60 min of sedation.

### Placing the tail vein line. Timing: 5 minutes

1. The tail vein should be clearly visible after 10 min on the heating pad. For mice over 30 g, this is typically sufficient for being able to place the IV line. For mice from 15–30 g, cover their heads with a paper towel so as not to damage their eyes, and heat them with a heating lamp 20 inches off the surface of the table for 3 min. At this point the veins will be fully dilated, and you can attempt to place the IV needle.
2. Set up the pump (rate (see **Table 2**), driveshaft distance, and direction) and syringe (on the side of the mat with the pump) before attempting to place the line.
3. Insert the IV needle into the tail vein near the tip of the tail.
4. To verify proper line placement, apply very gentle pressure to the end of the plunger to deliver 20–30 μL of the bolus dose (**Table 2**).
5. Once proper IV placement is confirmed, slowly push the remainder of the bolus by hand over ~30 sec. *Important:* If infusing acetate, push the bolus more slowly (over 60–90 sec) as injecting this tracer too quickly can cause respiratory arrest.
6. Once the bolus has been given, place the syringe onto syringe pump and begin infusion.

### Infusion. Timing: 2.5–6 hours

1. Infuse the tracer. Use Table 2 as a dosing guide.
2. Continuously monitor the mouse for signs of waking and patency of the infusion line.

### Terminal procedures. Timing: 5–10 minutes per mouse

*Note:* These procedures are designed for T cell isolation as a primary endpoint. If a different primary outcome is desired, it is recommended to collect and freeze that tissue/fluid as quickly as possible to minimize metabolomic artifacts from processing.

#### Blood collection

*Note:* This step is important to determine terminal enrichment of the labeled infused substrate in each mouse. For this purpose, whole blood, serum, or plasma is sufficient and can be used to accommodate the needs of the investigator.

1. At the end of tracer infusion, and while the mouse is still alive, lightly puncture the femoral vein with a 5/8” 27G needle (BD #305115) attached to a 1 mL syringe.
2. Transfer blood to a microcentrifuge tube and snap freeze for later processing.

#### Spleen collection for T cell isolation

1. Remove the spleen from the mouse and place in 10 mL of ice-cold HBSS.
2. Homogenize spleen in a 70 μm cell strainer using rubber end of 10 mL syringe.
3. Centrifuge homogenate at 500xg for 3 min at 4°C.
4. Use the resulting crude pellet for T cell isolation (see below).

#### Tissue/biofluid collection

Note: The goal of tissue collection is to quench metabolism as quickly as possible once it is removed from its native state.

1. Begin with the primary tissue/biofluid of interest. Remove the tissue/biofluid and fold into pre-labeled aluminum foil or place in a pre-labeled Eppendorf tube, and immediately freeze in liquid nitrogen. Note: Weighing fresh tissue before freezing not only complicates the workflow but also delays the quenching of metabolism. If tissue weight is desired, then it is recommended to weigh the frozen tissue at a later time. This is easier if tissues have been frozen in foil instead of tubes. Note: If tissues/biofluids are snap frozen in tubes, thermal expansion that occurs when the sample is moved from liquid nitrogen (−196°C) to a −80°C freezer can cause tubes to explode. Using a ~23G needle to puncture the caps of all tubes prior to the experiment will prevent this.
2. Store tissues/biofluids at −80°C for later processing.

### T cell isolation (Fig. 2). Timing: 35–45 minutes

**Figure 2.**
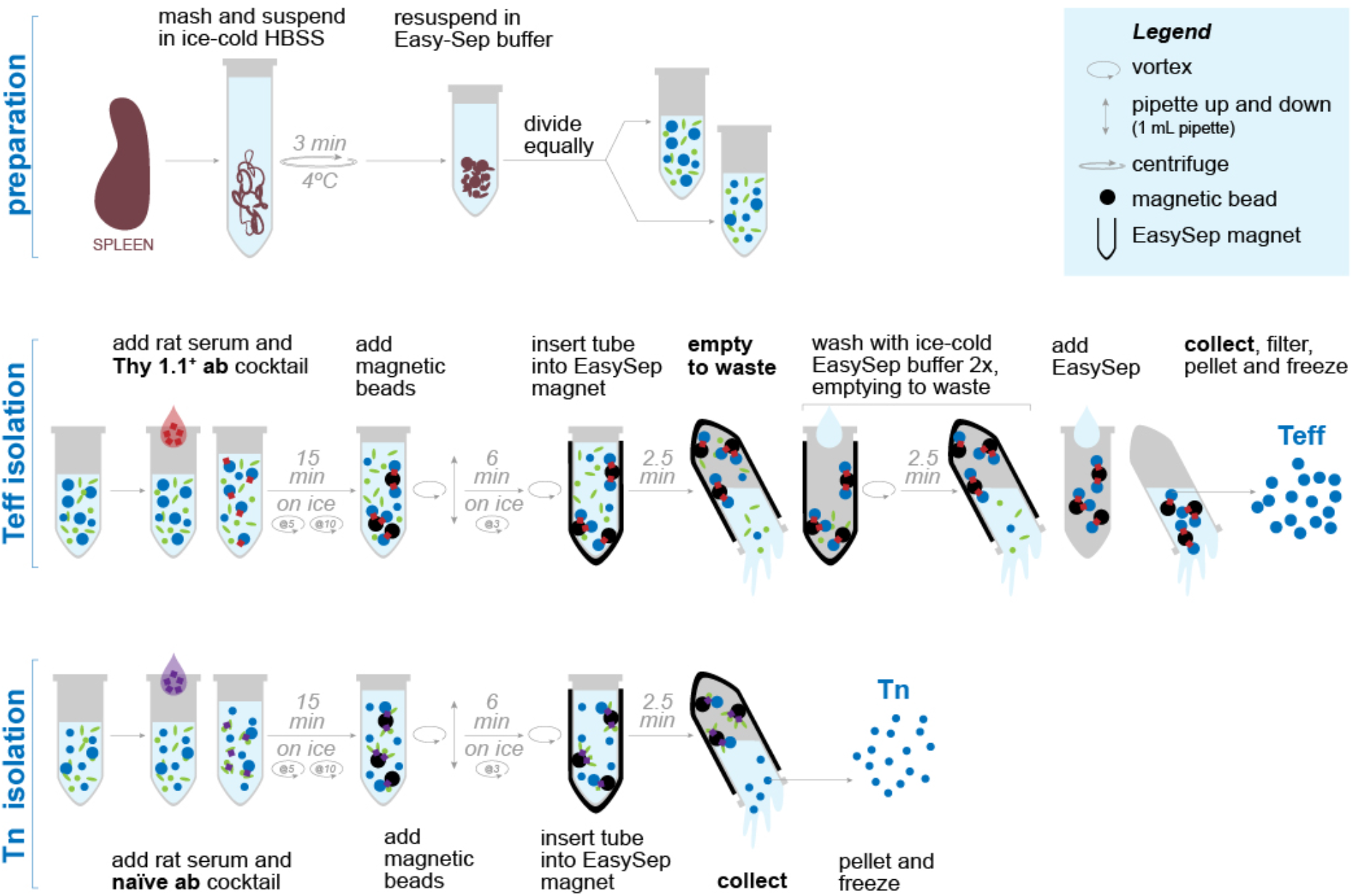
Bead isolation procedure. Small circular arrows with numbers enclosed (e.g., “@3”) designate the time points at which the tube should be vortexed. Blue dots represent T cells. Red or purple squares represent antibody cocktail that allows specific molecules to adhere to the magnetic beads.

Important: Proceed to this step immediately after spleen removal and homogenization.

1. Resuspend crude spleen homogenate pellet in 4 mL of Easy-Sep media.
2. Split the volume equally into two 15 mL polystyrene tubes, one for naïve CD8^+^ isolation and one for Thy1.1^+^CD8^+^ isolation. Note: This step is dependent on experimental question. If one is only interested in antigen-specific CD8^+^ activated T cells, there is no need to split volume into two tubes. Instead, resuspend the crude spleen pellet in 2 mL of Easy-Sep media (step 1). Important: Keep cells on ice for steps 3 and 4 below. Important: T cell pellets should be frozen within 35 minutes of removing the spleen.
3. For negative selection of naïve CD8^+^ cells:

a. Add 100 μL of rat serum to the 2 mL suspension, vortex for 5 seconds.
b. Add 100 μL of naïve antibody cocktail, vortex for 5 seconds.
c. Incubate for 15 min on ice, vortex for 5 seconds at the 5^th^ and 10^th^ minute.
d. Ensuring beads are completely resuspended, add 230 μL of beads and vortex for 5 seconds.
e. Incubate for 6 min on ice, vortex for 5 seconds at the 3^rd^ minute and at the end.
f. Insert the magnet for 2.5 min.
g. Collect the negative fraction by picking up the magnet and in a continuous motion invert the magnet with tube into a fresh falcon tube.
h. From the negative fraction, take 100–150 μL for cell counting and FACS analysis (below).
i. Centrifuge remaining cells in an Eppendorf tube at 500xg for 3 min at 4°C.
j. Aspirate supernatant carefully so as not to disturb the cell pellet. Dry the inside of the tube with a Kim-wipe.
k. Freeze the pellet on dry ice for later analysis.
4. For positive selection of Thy1.1^+^CD8^+^ cells:

a. Add 100 μL of rat serum to the 2 mL suspension, vortex for 5 seconds.
b. Add 50 μL of Thy1.1 antibody cocktail, vortex for 5 seconds.
c. Incubate for 15 min on ice, vortex for 5 seconds at the 5^th^ and 10^th^ minute.
d. Ensuring magnetic beads are completely resuspended, add 50 μL of beads vortex for 5 seconds.
e. Mix using a 1 mL pipette
f. Incubate for 6 min on ice, vortex (5 seconds) at the 3^rd^ minute and at the end.
g. Insert the magnet for 2.5 minutes.
h. Wash 2×

i. Discard the negative fraction from the tube by picking up the magnet and in a continuous motion invert the magnet with tube into a waste container. Dab off excess waste from the lip of the tube.
ii. Add 2 mL of fresh, ice-cold EasySep media.
iii. Take tube out of magnet and Vortex to resuspend beads on the sides of tube back into solution.
iv. Insert tube back into magnet and incubate for 2.5 minutes.
v. Repeat.
i. Dump the negative fraction from the tube into liquid waste.
j. Collect the positive fraction by adding 1 mL EasySep media into the tube.
k. Vortex to resuspend the positive fraction from the sides of the tube and filter through 70 μm cell strainer using P1000 micropipette into clean Eppendorf tube.
l. Of the filtrate, take 50–75 μL for cell counting and FACS analysis (below).
m. Centrifuge remaining cells in an Eppendorf tube at 500xg for 3 min at 4°C.
n. Aspirate supernatant carefully so as not to disturb the cell pellet. Dry the inside of the tube with a Kim-wipe.
o. Freeze the pellet on dry ice for later analysis.

### FACS. Timing: 2 hours

*Overview*: Analysis by flow cytometry is used to assess the quality of the T cell isolation procedure described above. These flow-sorted samples are not used for mass spectral analysis.

1. Place 50 μL of bead isolated Thy1.1^+^OT-I^+^ CD8^+^ into 96-well round bottom plate.
2. Add of 150 μL of flow wash (PBS containing 2% FBS, 0.02% NaN_3_) and centrifuge 500 g, 5 min, 4°C.
3. Flick out supernatant and flow stain samples (Table 3) using flow wash in the dark at 4°C for 1 hour in 50 μL.
4. Add 150 μL of flow wash and centrifuge 500 g, 5 minutes, 4 degrees.
5. Resuspend samples in 200 μL of flow wash.
6. Analyze cell populations using a flow cytometer (Beckan Coulter CytoFLEX or other compatible cytometer).

### Metabolite extraction. Timing: 1.5 hrs

#### Polar metabolite extraction from cells and blood

1. Prepare extraction solvent (80% methanol) and store at −20°C.
2. Extract metabolites

a. T cells: add 1 mL of extraction solvent to frozen cell pellets.
b. Blood: Add 960 μL of extraction solvent to a 40 μL aliquot of blood.
c. Process blanks: add extraction solvent to empty tubes and treat as biological samples through the entire workflow.
3. Vortex for 10 sec.
4. Incubate cells on dry ice for 1 hr to complete protein precipitation. Vortex.
5. Centrifuge extracts at max speed for 10 min to pellet proteins and nucleic acids.
6. Transfer 800 μL of supernatant to a fresh ultracentrifuge tube, being careful not to disrupt the pellet.
7. Take an additional 50–100 μL of supernatant from each sample and pool for a quality control sample (see **Box 2**).
8. Evaporate the extract to dryness in a Speedvac or under a gentle stream of N_2_ gas.

Pause point: Metabolite extracts can be stored dry at −80°C in a sealed tube for later analysis. As some metabolites, such as Acetyl-CoA and arginine, degrade more quickly it is recommended to complete MS analysis within one week.

### Metabolite extraction from tissues. Timing: 1.5 hours

*Note:* Here we describe the procedure for a Bligh-Dyer extraction^31^, which separates polar and non-polar metabolites into aqueous and organic layers, respectively. Compared to single-phase extractions, this method improves recovery and detection of nucleotides and other phosphate-containing compounds. If only amino acids and TCA cycle intermediates are desired, then extraction with 80% methanol is sufficient.

Important: In steps 1–7, it is essential to avoid sample thawing. It is recommended to practice the workflow before using analytical samples.

1. Pulverize tissues to a fine powder with a mortar and pestle equilibrated in liquid nitrogen. *Caution:* Mortar and pestle cause frostbite almost instantaneously. Use proper PPE. Tight fitting mechanics gloves underneath standard laboratory gloves are effective.
2. Transfer powdered tissue to a new, pre-chilled tube.
3. Label bead-mill homogenizer tubes. Equilibrate tubes and a weighing spatula in liquid nitrogen.
4. Quickly transfer chilled tube to scale and tare. While the scale is taring, open the sample tube and scoop sample with the chilled spatula but do not yet add to tube.
5. Once the scale is tared, add 20–30 mg of powdered tissue. *Important:* If this takes longer than 5–10 seconds to complete, return both sample tube and beadmill tube to liquid nitrogen and repeat.
6. Record the precise sample weight. This is best accomplished by a second person.
7. Immediately cap the weighed sample and return to liquid nitrogen.
8. Add 2:1 methanol:chloroform (1 mL per 40 mg tissue) to tissue samples and to an empty tube for a process blank. *Note:* Maintaining a consistent tissue:solvent ratio removes potential differences in extraction efficiency and normalized tissue weight/volume across all samples.
9. Homogenize using a bead-mill homogenizer for 30 seconds at maximum speed.
10. Add 1 part chloroform and 1.8 parts water. Vortex for 10 seconds.
11. Incubate on wet ice for 1 hour.
12. Centrifuge at maximum speed for 10 minutes.
13. Transfer tubes to wet ice, taking care not to disrupt separated phases.
14. The top (aqueous) phase contains polar metabolites including nucleotides, amino acids, sugar phosphates, and TCA cycle intermediates. The bottom (organic) layer contains non-polar metabolites and lipids. The interphase contains insoluble protein, nucleic acids, and other debris.
15. Without disturbing the organic or interphase, carefully transfer ~80% of the aqueous layer to a fresh tube. Take the same volume from every sample to ensure consistent tissue equivalents per vial.
16. Take an additional 50–100 μL of aqueous layer from each sample and pool for a quality control sample (see **Box 2**).
17. Evaporate the extract to dryness in a Speedvac or under a gentle stream of N_2_ gas.

Pause point: Metabolite extracts can be stored dry at −80°C in a sealed tube for later analysis. As some metabolites, such as Acetyl-CoA and arginine, degrade more quickly it is recommended to complete MS analysis within one week.

##### Box 2. Standards, blanks, and quality control samples for mass spectrometry metabolomics

Chromatography-mass spectrometry-based metabolomics relies on the separation of complex biological mixtures in time (column retention) and m/z (mass-to-charge) space. A single LC/MS run can yield over 25,000 features (a given m/z and a specific retention time), of which roughly ≤1,000 are true metabolites^32^. The remaining non-metabolite features are a combination of contaminants (e.g., leachables from plastics, dirty solvents, etc.), adducts (e.g., sodium), isotopes, and other artifacts. In addition, many metabolites have naturally occurring isomers (i.e., compounds with the same chemical formula but different structure/functions. Common examples include citrate/isocitrate, leucine/isoleucine, alanine/sarcosine, glucose/fructose, ATP/dGTP). Therefore, it is important to build controls into every metabolomics experiment to increase confidence in metabolite identification, identify false positives, and monitor instrument performance.

###### Double blanks

(typically only for LC/MS) The user will run the instrument (chromatography, scan, data collection) as if running a sample but without an injection. This allows for assessment of the cleanliness of the solvents and solvent path. Often, contaminants here are present through all or most of the analysis and can cause ion suppression, which can impede detection of low-abundance metabolites and their isotopes (an issue for stable isotope tracing studies).

###### Solvent blanks

These samples are neat solvent—used to resuspend the dried extracts for analysis—that are run through the method (chromatography, scan, data collection) as if they were a sample. In the present method, the solvent blank would be 50:50 (v/v) acetonitrile:H_2_O (LC/MS) or pyridine (GC/MS). Unlike the double blanks, solvent blanks pass through the injector loop and allow for identification of contaminants and carryover (compounds “stuck” to the injector).

###### Process blanks

These samples are created by adding extraction solvent directly to empty tubes and are treated identically to experimental samples throughout the entire workflow. These samples contain any contaminants introduced through the various processing steps and allow for identification of false-positive features in biological samples. These are arguably the most important blank, as all contaminant peaks present in double blanks and solvent blanks are contained here as well.

###### Pooled quality control (QC) samples

These are generated by pooling a small aliquot of each biological replicate into the same tube. As such, the sample matrix is identical to the samples being analyzed and each metabolite is present at a level intermediate to that of the samples being analyzed. These are critical controls that serve a number of important functions:

1. Initial column conditioning. Chromatography systems often require multiple (2–10) injections of sample at the beginning of every batch to stabilize compound retention and detection level.
2. Assessment of instrument performance. Depending on the number of samples and length of the chromatography, metabolomics studies can take hours to days to complete. During this time, it is important to be able to track shifts in instrument performance over time. This can be accomplished by running the pooled QC sample every ~10 injections throughout the study.
3. Control of time-dependent changes in analyte detection. Related to point 2 above, some analytes may increase or decrease as a function of time while waiting to be injected. Repeat pooled QC injections can allow for tracking and statistical correction of this effect if necessary. For discussion and tools for QC correction see reference by Yang *et al.* ^33^.

###### System suitability standards

These are identical samples that can be run at the beginning of every study to ensure optimal instrument performance. These are often small aliquots of pooled plasma/serum or a mixture of chemical standards. These allow for instrument performance tracking over months to years and can provide early detection of looming column failure or source contamination.

Below is an example template for setting up an LC/MS run.

Note: Randomize the order of experimental samples

Run 1. Double blank Run

Run 2. Solvent blank

Run 3. System suitability standard

Run 4. Process blank

Run 5–7. Repeat pooled QC injections for column conditioning. Additional QC injections can be added to ensure stability before moving to experimental samples. Ideally, you want a retention time difference <0.05 minutes and peak area difference <5% between the last two QC injections before beginning to run experimental samples.

Run 8–17. Randomized experimental samples 1–10

Run 18. Pooled QC

Run 19–n. Experimental samples

Run n+1. Pooled QC

Run n+2. Process blank

Run n+3. Double blank

#### Stable Isotope Labeling (SIL) analysis

Here we provide details for robust SIL analysis using both semi-targeted LC/MS and targeted GC/MS approaches. LC/MS is excellent for broad metabolite coverage, including nucleotides and sugar phosphates. Alternatively, GC/MS is well-suited to assess TCA cycle intermediates and most amino acids.

### Preparation of extracts for HILIC LC/MS analysis. Timing: 20 minutes

1. Resuspend extracts in 50:50 (v/v) acetonitrile:H_2_O. The resuspension volume should be determined *a priori* to yield desired sample concentration. Recommended volumes and on-column equivalents are list below.

a. T cells: 50 μL = 100,000 cells/μL; 200,000 cells/2 μL injection on column
b. Blood: 200 μL = 0.2 μL blood extract/μL; 0.4 μL/2 μL injection on column
c. Tissue: 100 μL = 160 μg/μL; 320 μg equivalents/2 μL injection on column
2. Vortex for 10 seconds.
3. Sonicate in a water bath sonicator for 5 minutes.
4. Vortex for 10 seconds.
5. Centrifuge at max speed for 1 min to pellet any insoluble metabolites and debris carry-over. Transfer supernatant to an autosampler vial with an insert. Note: A pellet may not be observable.
6. Maintain samples at 4°C until LC/MS analysis.

### LC/MS Analysis. Timing: 0.5–2 days

*Note:* The choice of column and solvent system should be based upon the desired metabolite coverage. Here, we present a basic pH, hydrophilic interaction (HILIC) separation that is excellent for separating and detecting a large number of polar compounds including nucleotides, sugar phosphates, amino acids, and carboxylates.

1. Prepare and condition a Sigma ZICpHILIC column with guard column according to manufacturer’s instructions.
2. Prepare mobile phases. *Note:* Fresh mobile phases should be made at least weekly and should not be added to remaining amounts of mobile phase (i.e. “topped off”). *Important:* always use LC/MS grade solvents and additives.

a. Mobile phase A: Water with 10mM ammonium acetate, 0.1% (v/v) ammonium hydroxide, and 0.1% (v/v) medronic acid (2.5 μM final). Note: Medronic acid, and also low levels (~10μM) of ammonium phosphate^34^, improves detection and chromatographic resolution of phosphate containing compounds.
b. Mobile phase B: 90% acetonitrile with 10mM ammonium acetate (770 mg/L), 0.1% (v/v) ammonium hydroxide, and 0.1% (v/v) medronic acid (2.5 μM final; Agilent, 5191-3940). *Note*: These salts can precipitate out of high organic solvent content. Dissolve ammonium acetate, ammonium hydroxide, and medronic acid in 100 mL of water. Then slowly add 900 mL of acetonitrile. If precipitation forms or the solvent appears cloudy, discard and restart. ***Failure to do so can damage the HPLC and column***.
c. Degas the solvents in a water batch sonicator for 10 minutes.
3. Turn on the mass spectrometer and allow ESI source voltage, gas flow, and temperature to equilibrate.

a. Capillary voltage (negative ion mode): 2500V
b. Sheath gas: 60 (arb)
c. Aux gas: 19 (arb)
d. Sweep gas: 1 (arb)
e. Ion transfer tube temperature: 300°C
f. Vaporizer temperature: 250°C
4. Purge each HPLC channel with mobile phase individually at 5 mL/minute for 5 minutes.
5. Ensure that column flow is diverted to waste (i.e., not going to the mass spectrometer).
6. Begin column flow (50% B) at 100 μL/minute for at least 20 minutes. If pressure is not stable (<5 bar fluctuation/minute), then purge the pumps again.
7. Switch to starting conditions (100%B, 150 μL/minute), and wait for pressure to stabilize (~5–10 min minutes).
8. Once pressure is stabilized and the ionization source is up to temperature, analysis can begin.
9. Inject 2 μL of sample on column.
10. Follow UHPLC analytical gradient in **Table 4**.
11. MS Scan parameters

**Table 4.**
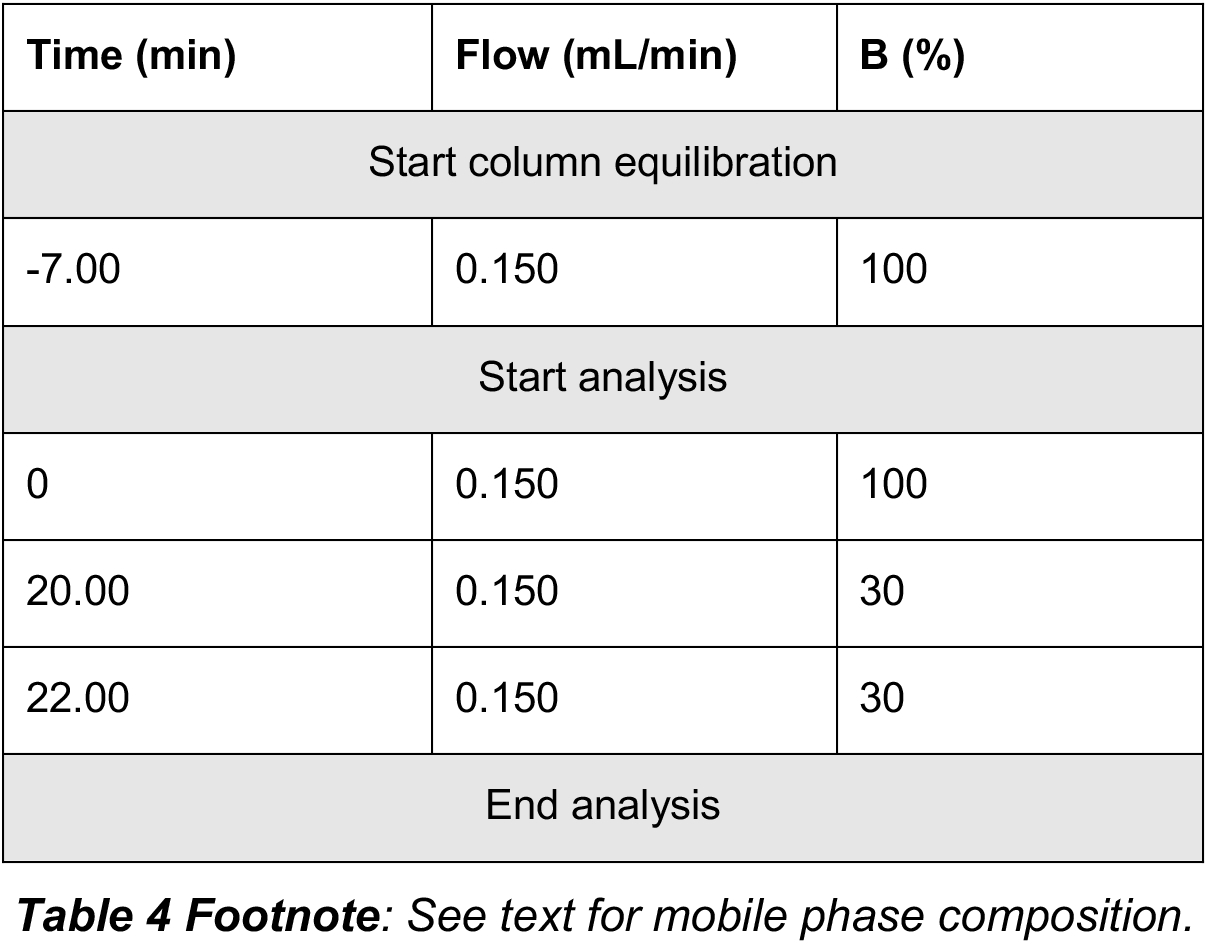
UHPLC analytical gradient schema.

Footnote: See text for mobile phase composition.
| Time (min) | Flow (mL/min) | B (%) |
| --- | --- | --- |
| Start column equilibration |  |  |
| -7.00 | 0.150 | 100 |
| Start analysis |  |  |
| 0 | 0.150 | 100 |
| 20.00 | 0.150 | 30 |
| 22.00 | 0.150 | 30 |
| End analysis |  |  |

*Note*: MS^2^ fragmentation is indispensable for compound identification, whereas MS^1^ scans are necessary for accurate and reliable peak integration and quantitation. However, gathering both MS^1^ and MS^2^ data in a single run decreases the number of MS^1^ scans performed and can negatively impact quantitation. To address this limitation, experimental replicates can be analyzed by purely MS^1^ scans, and pooled quality control samples can be analyzed using data-dependent MS^2^ (ddMS^2^). Thus, experimental replicates are analyzed with the maximum MS^1^ scans per peak, and compound identification is inferred from MS^2^ scans in pooled quality control samples.

a. MS^1^ scan – experimental samples:

i. Time: 0–22 min
ii. Orbitrap resolution: 240,000
iii. Scan range: 70–1000 m/z
iv. RF Lens: 60%
v. Ion polarity: negative
b. Data dependent MS^2^ (ddMS^2^) scan – pooled QC samples for identification only

i. Time: 0–22 min
ii. MS1 scan parameters: same as (a) above, but with 60,000 resolution
iii. ddMS^2^ parameters

- Intensity threshold = 2.0 × 10^4^
- High energy collisional dissociation (HCD) fragmentation scan:

- Stepped collision energy: 20%, 35%, 50%
- Orbitrap resolution: 30,000
- Maximum injection time: 0.054 seconds
- Collision-induced dissociation (CID) fragmentation scan:

- Fixed collision energy: 30%
- Orbitrap resolution: 30,000
- Maximum injection time: 0.054 seconds
iv. Total cycle time: 0.6 seconds

### Preparation of metabolite extracts for GC/MS analysis. Timing: 2 hours

*Note:* Metabolite derivatives may be prepared from samples previously used for LC/MS. To do this, dry the remaining extract from the LC/MS samples and proceed to step 1.

*Caution:* solvents used for derivatization should only be opened and used in a chemical fume hood. Always wear gloves when handling these solvents.

1. Dissolve methoxyamine HCl (MOX) in pyridine (10 mg/mL). Note: Pyridine is prone to oxidation and should be kept under N_2_ or other inert gas.
2. Using reverse pipetting, add 30 μL of MOX to each sample.
3. Vortex for 5 min.
4. Heat for 30 min at 70°C.
5. Using reverse pipetting, add 70 μL of MTBSFA + 1% TMCS
6. Pulse vortex.
7. Heat for 60 min at 70°C.
8. Allow samples to cool to room temperature before running on GC/MS.
9. Run all samples within 24 hours.

### GC/MS analysis. Timing: 0.5–2 days

1. Tune the mass spectrometer.
2. GC parameters (Agilent 7890):

a. Carrier gas: UHP Helium (99.995%)
b. Inlet is operated in split mode (10:1) at 250°C, septum purge flow = 3 mL/min, pressure = 8.2 psi.
c. Column flow = 1.0 mL/min.
d. Oven parameters:

i. Initial: 60°C for 1 min
ii. Ramp: from 60°C to 320°C at a rate of 10°C/min
iii. Hold: 320°C for 10 min.
3. Mass selective detector (MSD) parameters (Agilent 5977b):

a. Ionization: electron impact (−70 eV)
b. Quadrupole temperature: 180°C
c. Source temperature: 250°C
d. Scan parameters: range = 50–1000m/z; scan speed = 1562 u/second; frequency = 1.6 scans/second; cycle time = 630.57 milliseconds; step size = 0.1 m/z
e. Solvent delay: 9 minutes

### MS data analysis. Timing: 4–8 hours

#### Peak picking and integration

1. Identify and quantify metabolites using El-MAVEN^35^. Targeted compound lists with retention times for both LC/MS and GC/MS protocols described here are provided in **Table S1** and **Table S2**, respectively. *Important:* retention time for each analyte should be validated by neat chemical standards and compared against MS^2^ spectral matches in pooled QC samples. *Important:* for GC/MS analysis, the analyzed fragment must be a molecular ion (i.e., all carbons from the endogenous metabolite must be present in the fragment). Other fragments, called “qualifier ions”, that are not necessarily molecular ions can be used to verify compound identification.
2. Perform natural isotope correction using Labeled LCMS Workflow (Elucidata). *Note*: This requires the chemical formula of each compound as an input to correct for natural abundance. This can be challenging in GC/MS as the formula of the derivatized fragment is not always known or easily deduced. Formulas for derivatized fragments used for analysis of common metabolites are in **Table S2**. *Alternative*: Natural abundance correction can be performed using unlabeled control samples if chemical formula is unknown. In addition, this approach can help correct for experimental or instrument errors that cause a departure in isotopic distributions from theoretical values. However, this requires adding unlabeled control animals to the protocol and can significantly increase the size of the experiment. A tool for natural abundance correction using unlabeled samples can be found at http://fluxfix.science ^36^.

#### Normalization

We typically achieve 30–60% M+6 glucose labeling in the blood when using the glucose dosing schedule in **Table 1** (**Figure 3A–B**). To account for inter-infusion variability, it is important to normalize fractional enrichment data to the level of M+6 glucose observed in each animal.

**Figure 3.**
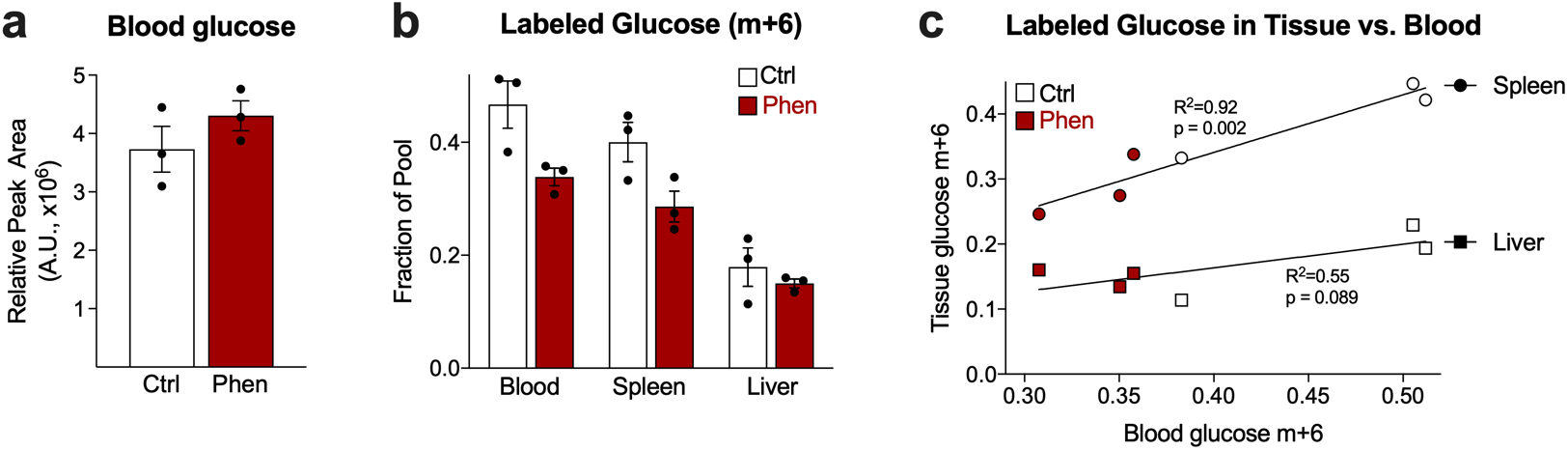
Blood glucose enrichment of ^13^C-glucose label in *Listeria*-infected mice. (**a**) Glucose levels in blood isolated from PBS- (Ctrl) or Phenformin- (Phen) treated *Listeria*-infected mice 2.5 dpi show no significant difference in blood glucose between treatment groups. (**b**) M+6 fractional enrichment of glucose in Ctrl and Phen treatment groups from (a). (**c**) Fraction of M+6 labeled glucose (of the total glucose pool) in spleen (circles) or liver (squares) relative to blood glucose M+6 enrichment in animals from (a). Blood glucose M+6 levels are predictive of spleen and liver M+6 glucose. Data represent mean ± SEM, with biological triplicates for each treatment group.

1. Generate a correction factor from M+6 glucose for each animal. To do this, divide the fraction of M+6 for each replicate by the average fraction of M+6 glucose for all samples.
2. Divide the fractional enrichment of each isotopologue by the correction factor for each sample.

#### Using total enrichment to visualize broad labeling patterns

SIL experiments can result in a large amount of data that can be difficult to visualize and interpret *en masse*. This is largely because each metabolite has a unique number of possible isotopic labeling patterns, resulting in a three-dimensional matrix (sample × metabolite × isotopologue). To address this, we reduce the dimensionality of the data set by calculating “total fractional enrichment.” This approach allows investigators to rapidly identify relevant metabolites on which to focus with more in-depth analysis.

1. Using the normalized isotope data (above), calculate the sum of all labeled isotopologues (i.e., M+1, M+2, …, M+n; do not include naturally occurring M+0) for each observation of metabolite. This should yield a simple a × b matrix, where a = biological replicate and b = metabolite.
2. Use this matrix of total fractional enrichment values to perform basic statistics and visualization tools, such as volcano plots and heatmaps.

### Timing

The dynamic nature of both metabolism and animal physiology in the acute post-infection period (<6 days) make timing and coordination of all phases of the experiment critical. Timing will depend on several factors: days post infection (dpi), length of fasting prior to infusion, infusion duration, duplicate equipment available (e.g., syringe pumps), and pharmacological treatment time. The beginning of the experiment (time = 0) is the infection of the mouse. The time of day and day of the week for infection should be selected so that infusions can occur during a regular workday. For example, in the experiment outlined in “Anticipated Results”, C57BL/6 mice were adoptively transferred with 2.5 × 10^6^ OT-I^+^Thy1.1^+^ CD8^+^ T cells on a Saturday, and then infected with 2 × 10^6^ CFU of attenuated LmOVA on the next day (Sunday) at 22:00h. This allowed the infusion for the 2.5 dpi timepoint to start on Wednesday at 10:00h. Six-hours prior (04:00h Wednesday) to the infusion, feed was removed from cages and the mice were treated with PBS or Phenformin (100 mg/kg) via IP injection. The mice were then given ^13^C-glucose infusion for 2.5 hours under isoflurane.

It is possible to infuse two mice per syringe pump. Multiple syringe pumps can accommodate even higher throughput while maintaining experimental timing. This is accomplished by staggering the beginning of infusions, such that mice on the second pump are initiated immediately after infusions begin on the first pump. Thus, the time difference between mice is ~20–30 minutes, as opposed to 3–6 hours (depending on the length of infusion) if only a single pump is used.

Another consideration is the time needed to isolate T cells from the mice (35–45 min) after the infusion is completed. If the infusion is done by just one person, staggering the infusion time between batches of mice allows for a more stable workflow. However, if multiple people are present, this may not be needed as each person can take on the role of isolating T cells from mice as soon as the infusion time point has been reached for a particular set of mice.

Once T cells are isolated and tissues collected, the samples can be stored at −80°C for long periods of time (up to a year or more) before mass spectrometry analysis. Post-storage, metabolite extraction and derivatization (for GC/MS) can be completed in as little as one day. GC/MS and LC/MS analysis duration is dependent on the number of samples and can take ~30–40 minutes per sample to run.

### Troubleshooting

#### Placing the tail vein needle

This step is critical for the success of this experiment. Below are troubleshooting tips.

1. If the line has been incorrectly placed, the insertion site will swell and turn white, and there will be resistance on the plunger.
2. If the needle is stuck in the interstitial space, it will be possible to push volume into the tail with heavy pressure, but doing so will cause the tail tissue to completely blanche.
3. If the needle has obstructed the vein, it will be possible to push the volume into the mouse, but you will see that all blood flushes out of the vein when pushing the bolus, and coloration does not return.
4. If any of these happen, the needle should be retracted and placement reattempted higher up the tail vein. You can try IV placement 3–4 times per side of the tail vein as long as each new attempt is “higher” up on the tail than the previous attempt (closer to the body of the mouse). This will avoid the solution leaking out of holes in the vein made from previous attempts once the IV is properly placed.
5. If the vein has collapsed, it is no longer usable.
6. If the tissue around the vein has blanched too much, you will be unable to see the vein to reattempt placement.
7. If the needle is inserted correctly:

b. The plunger of the needle will glide freely and without resistance when gentle pressure is applied.
c. The tail tissue will not whiten.
d. The vein will still be visible during and after the bolus push.

- *Note:* If the vein loses placement during the infusion, the area around the insertion site will whiten, and the needle should be removed and replaced.
- *Note:* If the needle maintains correct placement throughout the entire infusion, the mouse will bleed from the insertion site when the needle is removed at the end of the infusion.

### Anticipated Results

The protocol described here provides an effective and reproducible tool for achieving in vivo tracing. To demonstrate the power of this method, we characterized in vivo glucose utilization in response to acute phenformin treatment in mice 2.5 dpi by *LmOVA*. Phenformin (Phen) is a biguanide that inhibits complex I of the respiratory chain and ultimately can result in TCA cycle collapse and reduced oxidative phosphorylation (OXPHOS)^37^. Mice were infused with U-^13^Cglucose (as in **Table 2**) 6-hours post Phen injection. Following tracer infusion, blood glucose was not significantly different between groups (**Figure 3A**). Total blood enrichment of M+6 glucose was between 30–50% (**Figure 3B**), which is typical variability when following the glucose dosing schedule in **Table 2**. To account for variability, we normalized mass isotopologue distributions (MID; see note on nomenclature in **Box 1**) of each metabolite to glucose M+6. Blood glucose levels strongly but imperfectly correlate with tissue M+6 levels (**Figure 3C**); thus, we prefer to normalize to glucose M+6 fraction in the tissue of interest to best capture the local milieu. In the case of T cells, we normalized to spleen glucose M+6 fraction.

Flow cytometric analysis of cells post-isolation provided us with an idea of the cell population (homogenous/heterogenous) we were analyzing. The magnetic bead-based T cell sorting procedure we describe here resulted in a highly pure and viable population of Thy1.1^+^ cells, with the vast majority of the Thy1.1^+^ cells classified as early effector cells (EECs) (KLRG1^−^IL-7R^−^) in both PBS (Ctrl) and Phen treated mice at 2.5 dpi (**Figure 4**). We traced ^13^C glucose into the TCA cycle in Teff cells isolated from PBS- and Phen-treated animals (**Figure 5A**). Calculating the total enrichment (i.e. the sum of all labeled isotopologues) of each metabolite present revealed 21 differentially ^13^C-enriched metabolites between Ctrl- and Phen-injected mice. Visualization oh these data in a heatmap revealed a that Phen induced a rerouting of glucose metabolism in ex vivo isolated OT1^+^ T cells (Figure 5B). On the whole, Phen promoted the conversion of glucose-dependent carbon through glycolysis and into lactate, alanine, and the TCA cycle. Interestingly, though Phen decreased TCA cycle intermediate abundance in transformed lymphocytes^37^, our data indicates the fraction of the TCA cycle derived from glucose in T cells is maintained in Phen-treated animals (**Figure 5A**). We further observed a broad decrease in ^13^C-glucose labeling in pyrimidine (e.g., UMP/UDP/UTP, CMP/dCMP, dCDP) and purine (e.g., ADP, GTP) nucleotides and phospholipid precursors (e.g., CDP-choline, CDP-ethanolamine) (**Figure 5B**). We hypothesize that this increased glycolytic disposal of glucose in response to Phen treatment may serve to compensate for impaired mitochondrial oxidative ATP production. Consequently, this pulls carbon away from both the pentose phosphate pathway and glutamate cataplerosis, both of which are necessary to support de novo nucleotide synthesis (**Figure 6**). Thus, these data suggest that Phen treatment in vivo promotes glycolytic disposal of glucose over nucleotide biosynthesis in activated T cells.

**Figure 4.**
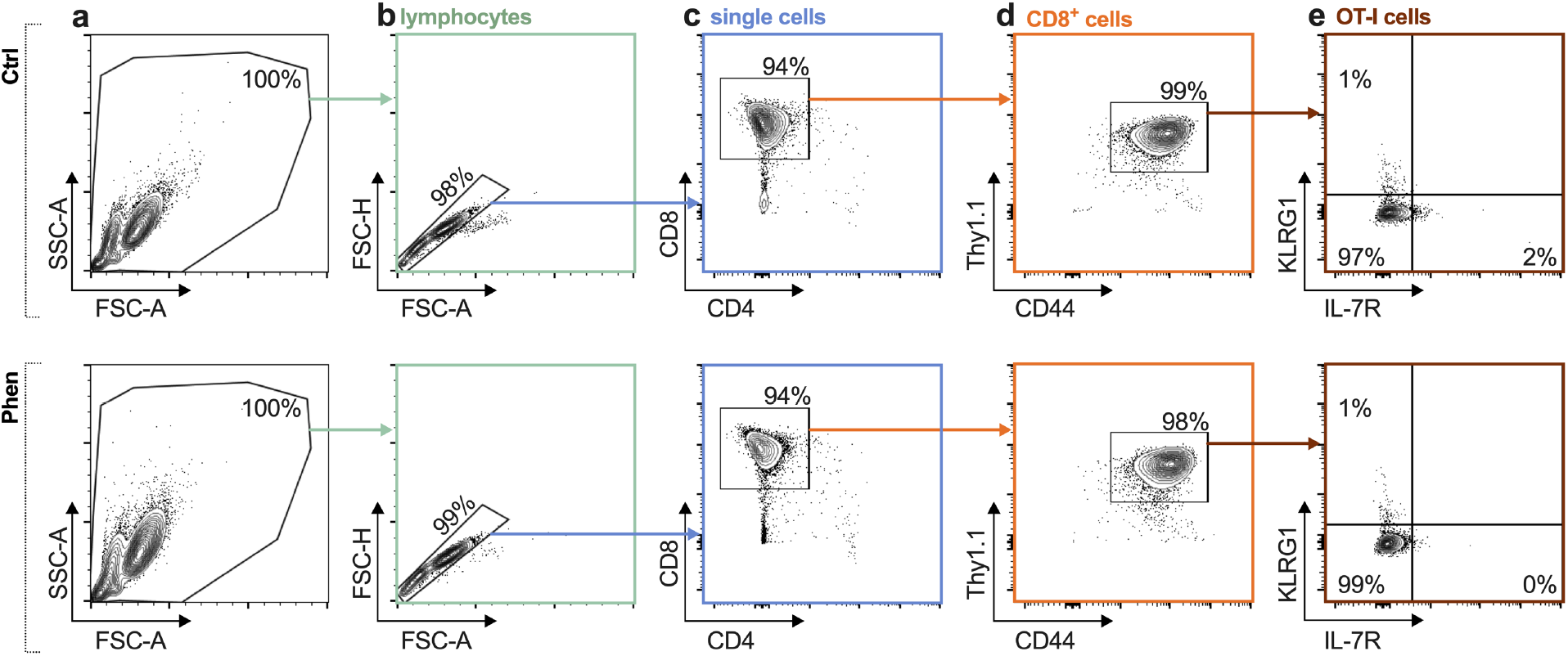
FACS analysis of T cells following magnetic bead isolation procedure. Shown are representative flow cytometry plots of T cells isolated from spleens of PBS- (Ctrl) or Phenformin- (Phen) treated *Listeria*-infected mice 2.5 dpi. The panels display: (**a**) FSC versus SSC parameters to identify lymphocyte gates; (**b**) FSC height versus area to identify single cells; (**c**) CD8 versus CD4 gating to identify CD8+ T cells; (**d**) Thy1.1 versus CD44 profiles to identify activated adoptively transferred OT-I T cells; and (**e**) KLRG1 versus IL-7R (CD127) staining for phenotypic characterization of Thy1.1^+^OT-I^+^ CD8^+^ T cells. Numbers indicate the percentage of cells in each gated region.

**Figure 5.**
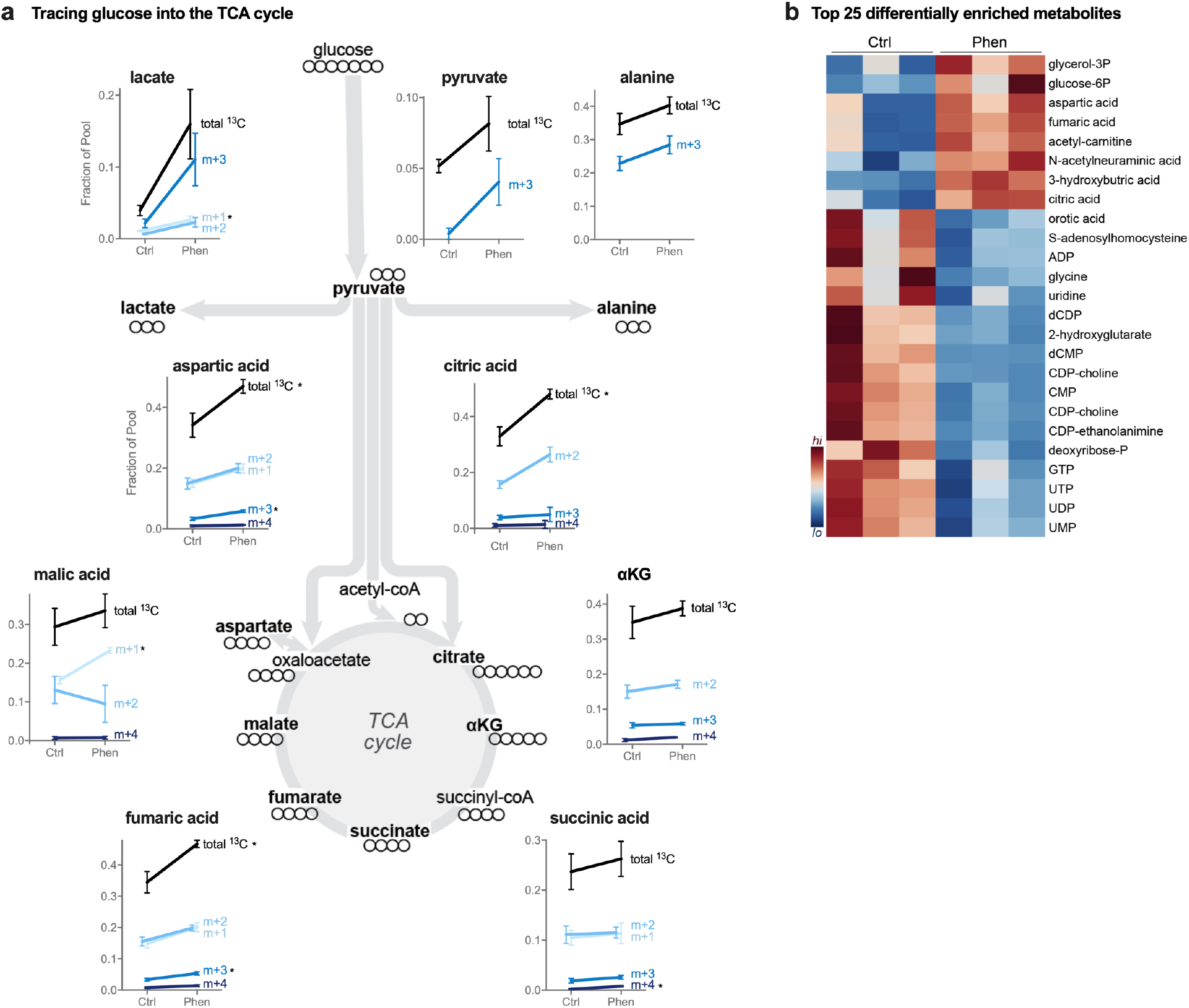
^13^C-glucose labeling patterns in T cells isolated from *Listeria*-infected mice. (**a**) MIDs of ^13^C-glucose-derived glycolytic and TCA cycle intermediates in OT-I T cells isolated from PBS- (Ctrl) or Phenformin- (Phen)treated *Listeria*-infected mice 2.5 dpi. Shown is the fraction of total ^13^C labeling (black) and individual MIDs (blue) for indicated metabolites relative to the total metabolite pool. Significant changes in MID patterns between Ctrl and Phen conditions are indicated by * (*p*<0.05). Phen increases the fraction of lactate, pyruvate, alanine, and TCA cycle intermediates derived from glucose. (**b**) Total enrichment of ^13^C-glucose-derived metabolites in T cells from Phenformin-treated animals. Shown is a heatmap of top differentially enriched metabolites. Data represent mean ± SEM, with biological triplicates for each treatment group.

**Figure 6.**
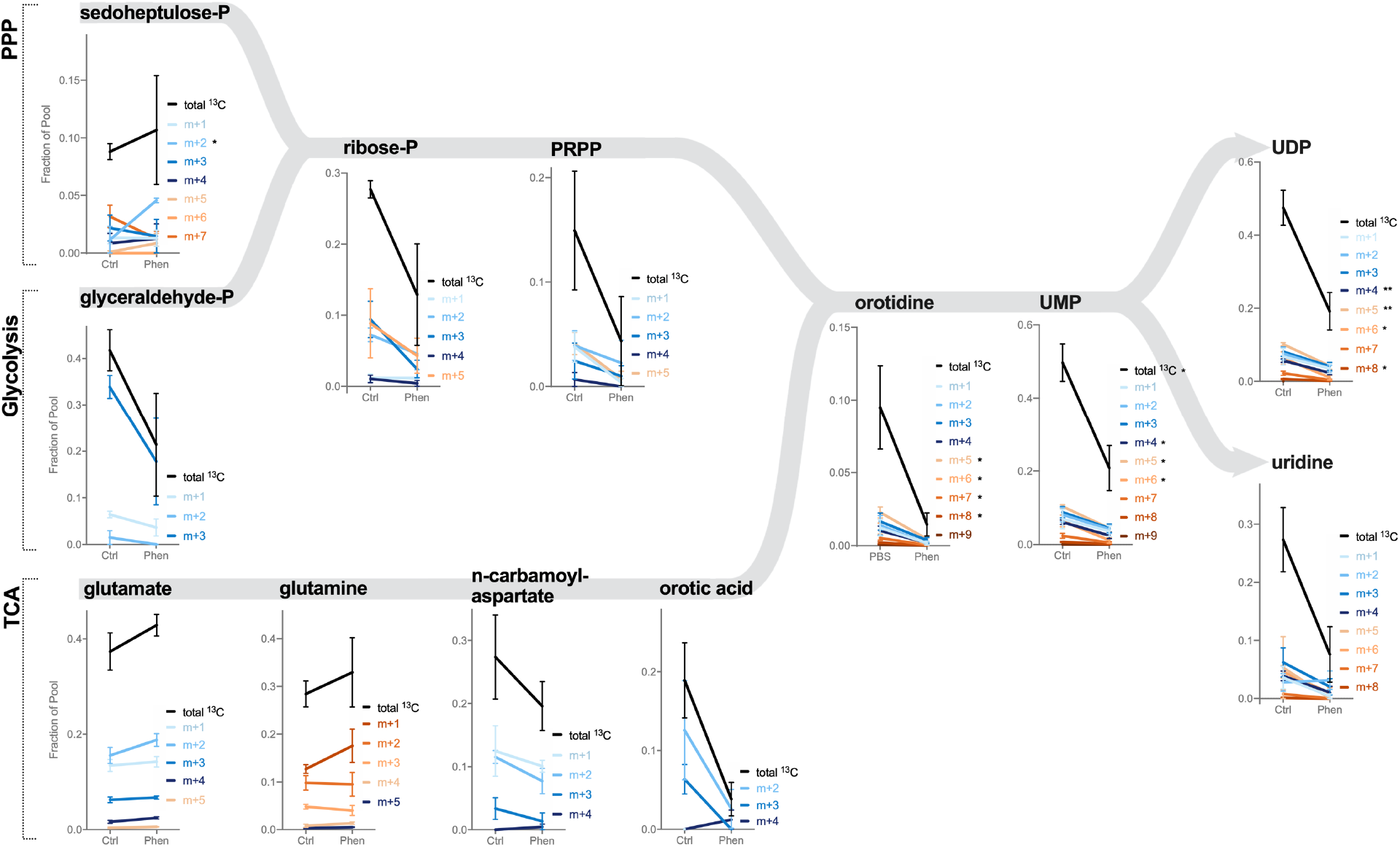
Pathway analysis of ^13^C-glucose-depedent nucleotide metabolism in activated T cells from *Listeria*-infected mice. Shown are MIDs of ^13^C-glucose-derived metabolites from the pentose phosphate pathway (PPP) and TCA cycle contributing to pyrimidine biosynthesis in OT-I T cells isolated from PBS (Ctrl) or Phenformin-treated *Listeria*-infected mice 2.5 dpi. Shown is the fractional enrichment of total ^13^C label (black) and individual MIDs (in color) for indicated metabolites. Significant changes in MID patterns between Ctrl and Phen conditions are indicated by * (*p*<0.05). Phen decreases glucose contribution to pyrimidine biosynthesis through both arms of the biosynthetic pathway. Data represent mean ± SEM, with biological triplicates for each treatment group.

## Supplementary Information

**Figure S1.** Preparation of infusion lines

**Table S1.** Targeted HILIC LC/MS compound list.

**Table S2.** GC/MS compound list.

## Author Contributions

Conceptualization, RDS, EHM, RGJ; Validation, RDS and EHM; Investigation, RDS, LMD, and EHM; Data Curation, RDS, EHM; Writing – Original Draft, RDS and EHM; Writing – Review & Editing, RDS, EHM, KSW, and RGJ; Visualization, KSW; Supervision, RGJ; Funding Acquisition, RGJ.

## Acknowledgements

We thank Elissa Levine, Rachel Sheridan, and staff from the Van Andel Institute (VAI) Metabolomics and Bioenergetics and Flow Cytometry core facilities for technical assistance. We thank Jeanie Wedberg and Michelle Minard for administrative assistance.

## References

1. Cham, C. M., Driessens, G., O’Keefe, J. P. & Gajewski, T. F. Glucose deprivation inhibits multiple key gene expression events and effector functions in CD8+ T cells. Eur. J. Immunol. 38, 2438–2450 (2008).

2. Chang, C. H. et al. Posttranscriptional control of T cell effector function by aerobic glycolysis. Cell 153, 1239 (2013).

3. Blagih, J. et al. The energy sensor AMPK regulates T cell metabolic adaptation and effector responses in vivo. Immunity 42, 41–54 (2015).

4. Balmer, M. L. et al. Memory CD8+ T Cells Require Increased Concentrations of Acetate Induced by Stress for Optimal Function. Immunity 44, 1312–1324 (2016).

5. Ma, E. H. et al. Serine Is an Essential Metabolite for Effector T Cell Expansion. Cell Metab. 25, 345–357 (2017).

6. Davidson, S. M. et al. Environment Impacts the Metabolic Dependencies of Ras-Driven Non-Small Cell Lung Cancer. Cell Metab. 23, 517–28 (2016).

7. Cantor, J. R. The Rise of Physiologic Media. Trends in Cell Biology 29, 854–861 (2019).

8. Ackermann, T. & Tardito, S. Cell culture medium formulation and its implications in cancer metabolism. Trends in Cancer 5, 329–332 (2019).

9. Vande Voorde, J. et al. Improving the metabolic fidelity of cancer models with a physiological cell culture medium. Sci. Adv. 5, eaau7314 (2019).

10. Cantor, J. R. et al. Physiologic medium rewires cellular metabolism and reveals uric acid as an endogenous inhibitor of UMP synthase HHS Public Access. Cell 169, 258–272 (2017).

11. Hensley, C. T. et al. Metabolic Heterogeneity in Human Lung Tumors. Cell 164, 681–694 (2016).

12. Sellers, K. et al. Pyruvate carboxylase is critical for non-small-cell lung cancer proliferation. J. Clin. Invest. 125, 687–698 (2015).

13. Faubert, B. et al. Lactate Metabolism in Human Lung Tumors. Cell 171, 358–371.e9 (2017).

14. Yuan, M. et al. Ex vivo and in vivo stable isotope labelling of central carbon metabolism and related pathways with analysis by LC–MS/MS. Nat. Protoc. (2019). doi:10.1038/s41596-018-0102-x

15. Llufrio, E. M., Wang, L., Naser, F. J. & Patti, G. J. Sorting cells alters their redox state and cellular metabolome. Redox Biol. 16, 381–387 (2018).

16. Binek, A. et al. Flow Cytometry Has a Significant Impact on the Cellular Metabolome. J. Proteome Res. 18, 169–181 (2019).

17. Chen, W. W., Freinkman, E. & Sabatini, D. M. Rapid immunopurification of mitochondria for metabolite profiling and absolute quantification of matrix metabolites. Nat. Protoc. 12, 2215–2231 (2017).

18. Ma, E. H. et al. Metabolic Profiling Using Stable Isotope Tracing Reveals Distinct Patterns of Glucose Utilization by Physiologically Activated CD8+ T Cells. Immunity (2019). doi:10.1016/J.IMMUNI.2019.09.003

19. Dyar, K. A. et al. Atlas of Circadian Metabolism Reveals System-wide Coordination and Communication between Clocks. Cell 174, 1571–1585.e11 (2018).

20. Eckel-Mahan, K. L. et al. Reprogramming of the circadian clock by nutritional challenge. Cell 155, 1464–1478 (2013).

21. Kinouchi, K. et al. Fasting Imparts a Switch to Alternative Daily Pathways in Liver and Muscle. Cell Rep. 25, 3299–3314.e6 (2018).

22. Overmyer, K. A., Thonusin, C., Qi, N. R., Burant, C. F. & Evans, C. R. Impact of anesthesia and euthanasia on metabolomics of mammalian tissues: Studies in a C57BL/6J mouse model. PLoS One 10, (2015).

23. Guo, N. L., Zhang, J. X., Wu, J. P. & Xu, Y. H. Isoflurane promotes glucose metabolism through up-regulation of miR-21 and suppresses mitochondrial oxidative phosphorylation in ovarian scancer cells. Biosci. Rep. 37, (2017).

24. Rao, L. K., Flaker, A. M., Friedel, C. C. & Kharasch, E. D. Role of Cytochrome P4502B6 Polymorphisms in Ketamine Metabolism and Clearance. Anesthesiology 125, 1103–1112 (2016).

25. Kharasch, E. D. & Thummel, K. E. Identification of cytochrome P450 2E1 as the predominant enzyme catalyzing human liver microsomal defluorination of sevoflurane, isoflurane, and methoxyflurane. Anesthesiology 79, 795–807 (1993).

26. Best, J. A. et al. Transcriptional insights into the CD8 + T cell response to infection and memory T cell formation. Nat. Immunol. (2013). doi:10.1038/ni.2536

27. Wang, Y. et al. Uncoupling Hepatic Oxidative Phosphorylation Reduces Tumor Growth in Two Murine Models of Colon Cancer. Cell Rep. (2018). doi:10.1016/j.celrep.2018.06.008

28. Hui, S. et al. Glucose feeds the TCA cycle via circulating lactate. Nature 551, 115–118 (2017).

29. Tardito, S. et al. Glutamine synthetase activity fuels nucleotide biosynthesis and supports growth of glutamine-restricted glioblastoma. Nat. Cell Biol. (2015). doi:10.1038/ncb3272

30. Marin-Valencia, I. et al. Analysis of tumor metabolism reveals mitochondrial glucose oxidation in genetically diverse human glioblastomas in the mouse brain in vivo. Cell Metab. (2012). doi:10.1016/j.cmet.2012.05.001

31. Bligh, E. G. & Dyer, W. J. A rapid method of total lipid extraction and purification. Can. J. Biochem. Physiol. 37, 911–917 (1959).

32. Mahieu, N. G. & Patti, G. J. Systems-Level Annotation of a Metabolomics Data Set Reduces 25 000 Features to Fewer than 1000 Unique Metabolites. Anal. Chem. 89, 10397–10406 (2017).

33. Yang, Q. et al. NOREVA: enhanced normalization and evaluation of time-course and multi-class metabolomic data. Nucleic Acids Res. (2020). doi:10.1093/nar/gkaa258

34. Spalding, J. L., Naser, F. J., Mahieu, N. G., Johnson, S. L. & Patti, G. J. Trace Phosphate Improves ZIC-pHILIC Peak Shape, Sensitivity, and Coverage for Untargeted Metabolomics. J. Proteome Res. (2018). doi:10.1021/acs.jproteome.8b00487

35. Agrawal, S. et al. EL-MAVEN: A fast, robust, and user-friendly mass spectrometry data processing engine for metabolomics. in Methods in Molecular Biology 1978, 301–321 (Humana Press Inc., 2019).

36. Trefely, S., Ashwell, P. & Snyder, N. W. FluxFix: Automatic isotopologue normalization for metabolic tracer analysis. BMC Bioinformatics 17, 1–8 (2016).

37. Izreig, S. et al. Repression of LKB1 by miR-17∼92 sensitizes MYC-dependent lymphoma to biguanide treatment. bioRxiv 2019.12.20.883025 (2019). doi:10.1101/2019.12.20.883025

