## Supplemental Figures for "In vivo metabolite tracing of T cells"

### Supplemental Materials for:
### In vivo metabolite tracing of T cells

Ryan D. Sheldon*, Eric H. Ma*, Lisa M. DeCamp, Kelsey S. Williams, Russell G. Jones

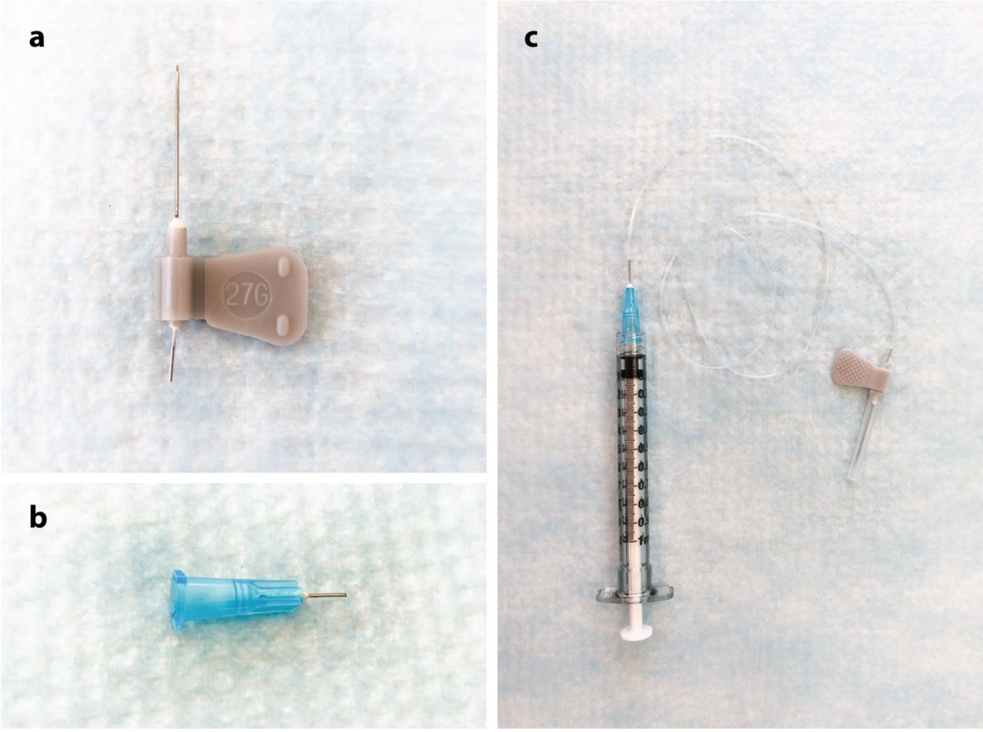


**Figure S1**. **Preparation of infusion lines.**

(**a**) 27G scalp vein assembly with line and one butterfly wing removed.

(**b**) 1.5” 25G needle cut to 0.5” in length.

(**c**) The needle from (b) is attached to the butterfly needle in (a) via 18”–20” of tubing (0.033” outer diameter, 0.014” inner diameter; MRE 033, Braintree Scientific). Please see main text for details.

**Table S1. Targeted HILIC LC/MS compound list.** Compound list using the described HILIC LC/MS method. *Important:* Retention times will vary from lab to lab and should be confirmed by chemical standards.

**Table S2. GC/MS compound list**. Compound list using the described GC/MS method. The quantifier ion is a molecular ion that can be used for isotope analysis. The qualifier ion should have the same retention and is used to improve confidence in compound identification. *Important:* Retention times will vary from lab to lab and should be confirmed by chemical standards.
